# Identification and comparison of *N*-glycome profiles from common dietary protein ingredients

**DOI:** 10.1101/2024.09.17.613496

**Authors:** Matthew Bolino, İzzet Avcı, Hacı Mehmet Kayili, Hatice Duman, Bekir Salih, Sercan Karav, Steven A. Frese

**Affiliations:** Department of Nutrition, University of Nevada, Reno; Reno, NV USA 89557; Department of Chemistry, Faculty of Science; Hacettepe University, 06500 Ankara, TR; Department of Biomedical Engineering, Faculty of Engineering; Karabük University; 78000 Karabük, TR; Department of Molecular Biology and Genetics, Çanakkale Onsekiz Mart University; 17020 Çanakkale, TR; University of Nevada, Reno School of Medicine; Reno, NV USA 89557

## Abstract

The *N-*glycome profiles purified from dietary bovine whey, egg white, pea, soy protein isolates and a recently commercialized animal-free whey is described. Purified glycoproteins resulting from centrifugation and ethanol precipitation of protein powder supplements were treated with peptide-*N-* glycosidase F (PNGase F) to release protein-bound *N*-glycans. Once released from the protein, *N*-glycans were labeled by procainamide labeling, purified via cotton-hydrophilic interaction liquid chromatography (HILIC), and analyzed using HILIC high performance liquid chromatography equipped with a fluorescence detector and a quadrupole time-of-flight tandem mass spectrometry (HILIC-FLD-QTOF-MS/MS). A total of 33, 33, 10, and 10 *N-*glycan structures were identified from bovine whey, egg, soy, and pea glycoprotein isolates, respectively. The type of *N-*glycans per glycoprotein source were highly predictable, likely attributed to differences in biosynthetic glycosylation pathways. Mammalian glycoprotein sources favored a combination of complex and hybrid glycan configurations while the plant proteins were dominated by oligomannosidic *N*-glycans. Bovine whey glycoprotein isolate contained the most diverse *N-*glycans by monosaccharide composition as well as structure, while plant sources such as pea and soy glycoprotein isolates contained an overlap of oligomannosidic *N-*glycans. The results suggest *N-*glycan structure and composition is dependent on the host organism rather than protein sequence homology, likely driven by the differences in *N-*glycan biosynthetic pathways.

## 1. Introduction

*N-*linked glycans are complex carbohydrate moieties bound to asparagine residues of many cellular proteins through a process known as glycosylation (Fernández-Tejada et al., 2015). Glycosylation is one of several post-translational modifications to protein structures, and glycosylation can be responsible for protein function as well as the stability, solubility, and structure of proteins (Molinari, 2007; Skropeta, 2009; Stanley et al., 2015). During *N*-glycan synthesis, a 14 subunit *N-*glycan (Glc3Man9GlcNAc2) synthesized in the endoplasmic reticulum (ER) is transferred from the lipid anchor, dolichol pyrophosphate, to an asparagine residue linked via an *N*-acetylglucosamine within a specific *N*-glycosylation acceptor sequence (Asn-X-Ser/Thr) of the recipient protein (Bieberich, 2014). The *N*-glycan structure is then modified and “trimmed” in the ER and Golgi apparatus by hydrolytic removal of sugar residues followed by re-glycosylation or “processing” by the addition of new monomers such as galactose, fucose, or mannose, and the composition and architecture of the resulting *N*-glycan is dependent on the organism and the particular glycosylation site and/or glycoprotein (Bieberich, 2014). *N-*linked glycans act as a quality control checkpoint for proper protein folding in the ER, resulting in the export of the protein from the ER or tagging the protein for degradation (Aebi et al., 2010; Helenius & Aebi, 2004). Additionally, other cellular roles such as protein transport, migration, and adhesion have also been attributed to glycosylation (Bieberich, 2014).

*N-*glycans found within dietary glycoproteins or derived from the host may also serve as energy substrates for the adult microbiota, especially when fiber intake is low. N-glycoproteins ingested from diet or shed host epithelial cells are likely the primary sources of *N*-glycans (Koropatkin et al., 2012). Among infants, *N-*glycans bound to human milk proteins can serve as important substrates for an infant gut microbe, *Bifidobacterium longum* subsp. *infantis* (*B. infantis*; Barratt et al., 2022; Karav et al., 2019; Karav, Parc, et al., 2015). For example, *B. infantis* has been shown to release *N-*glycans from human milk proteins *in vivo* (Karav et al., 2019) and access to available *N-*glycans can serve as an important fitness determinant for *B. infantis* (Barratt et al., 2022). There is also evidence that *N*-glycans can serve as prebiotics (Barratt et al., 2022; Karav et al., 2016). However, most of the knowledge of the *N*-glycome of dietary protein sources has focused on bovine and human milk (Barboza et al., 2012; Dallas et al., 2011; Karav, Bell, et al., 2015; Nwosu et al., 2012; Parc et al., 2015; Smilowitz et al., 2013; Zivkovic et al., 2011), and much of the knowledge of *N*-glycan utilization by the gut microbiome has focused on individual constituent microbes (Briliūtė et al., 2019; Crouch et al., 2022).

Structural differences in *N-*glycomes between human and other mammalian milk proteins are well characterized (Barboza et al., 2012; Nwosu et al., 2012; Smilowitz et al., 2013; Zivkovic et al., 2011), but other areas of research have characterized the biosynthetic systems among a wide variety of organisms, demonstrating an incredible array of potential *N*-glycan structures arising from variations in *N*-glycan biosynthetic capabilities and regulatory networks (Stanley et al., 2015). For example, mammalian bovine milk proteins contain complex, hybrid, and oligomannose *N-*glycans (Nwosu et al., 2012) while plant protein glycosylation is primarily described as oligomannosidic *N-*glycans, with no hybrid or complex types present and the inclusion of distinct carbohydrate monomers, such as arabinose and xylose (Castilho et al., 2011; Strasser, 2016). How differences in *N-*glycan structure and composition impact the composition and function of the gut microbiome is not understood, and there is currently a paucity of knowledge as to the structural composition of dietary *N*-glycans among common sources in the diet to address this gap in knowledge. This study sought to characterize the *N*-glycan structures from four glycoproteins sources that are widely consumed in whole and processed foods; egg white (from *Gallus gallus domesticus*), bovine whey protein (from *Bos taurus*), pea (from *Pisum sativum)*, and soybean (from *Glycine max*).

## 2. Methods

### 2.1 Protein Purification

Protein was purified from commercially available protein isolates derived from large-scale commodity ingredient processing in the United States. Each sample was subjected to four rounds of ethanol precipitation by adding four volumes of ice-cold ethanol, incubation at -2^°^C overnight, then followed by centrifugation at 4°C (4000RPM, 25 mins) to remove residual sugars and other remaining contaminants. The protein samples were subsequently aliquoted and dried at 30°C under vacuum centrifugation (Eppendorf 5301 Vacufuge Concentrator System). After rehydration with distilled H_2_O, the purified protein was then quantified using a Qubit BR Protein assay (ThermoFisher Scientific, Waltham, MA USA).

### 2.2 N-glycan deglycosylation

Initially, 0.5 mg of each sample was transferred into microcentrifuge tubes and 50 µl of 1% SDS was added to each sample to facilitate protein solubilization. Samples were then subjected to incubation at 70°C for 10 minutes. Following this, 25 µl of 4% NP-40 and 25 µl of 5X PBS were added and the mixture was gently vortexed to ensure thorough mixing. Subsequently, 1U of PNGase F obtained from Promega (Madison, WI, USA) was added to each sample to enzymatically release *N*-linked glycans from the glycoprotein substrates and incubated overnight at 37°C.

### 2.3 N-glycan labeling

To label released *N*-glycans, 50 µl of the procainamide labeling (110 mg/ml procainamide in a 10:3 (v/v) mixture of DMSO and glacial acetic acid) and 50 µl of the NaCNBH_3_ (63 mg/ml NaCNBH_3_ in a 10:3 (v/v) mixture of DMSO and glacial acetic acid) were added to the samples and incubated at 65°C for 2 hours. After the incubation period, samples were centrifuged at maximum speed for 5 minutes to pellet any insoluble materials. The supernatant containing the labeled *N-*glycans was carefully transferred to fresh microcentrifuge tubes for subsequent analysis.

### 2.4 N-glycan purification

To purify *N*-glycans, cotton-HILIC was used as described previously (Kayili & Salih, 2021). Briefly, a cotton wool plug was inserted into a pipette tip (100 μL capacity). The cotton wool-containing pipette tip underwent a washing procedure consisting of three rinses with pure water followed by three rinses with an 85% acetonitrile (ACN) solution. Following this, a loading solution was prepared by mixing 15 μL of procainamide-labeled *N*-glycan sample with 85 μL of ACN. Each sample (loading solution) was then aspirated and dispensed 15 times using a cotton wool-containing pipette tip. Subsequently, each cotton wool-containing pipette tip underwent a washing process comprising of five rinses with 100 μL of a solution containing 85% ACN, 14% water, and 1% trifluoroacetic acid (TFA) (v/v/v), followed by five rinses with an 85/15 ACN/water (v/v) solution. Finally, the *N*-glycans that were loaded onto the cotton wool were eluted by aspirating and dispensing 10 times with 25 μL of pure water.

### 2.5 HPLC-HILIC-FLD-QTOF-MS/MS analysis of N-glycans

Analysis of procainamide-labeled *N*-glycans was performed as described (Kayili, 2020) on a QTOF (TIMSTOF) mass spectrometer (Bruker Daltonik, GmbH) coupled with an Agilent 1200 series HPLC system featuring a fluorescence detector. Separation of the labeled *N*-glycans were achieved with a Waters Glycan BEH Amide column (2.5 μm, 2.1 mm ID x 15 cm L). The fluorescence detector was set with excitation and emission wavelengths of 310 nm and 370 nm, respectively. Mobile phases consisted of 100% acetonitrile (A) and 50 mM ammonium formate (pH: 4.4) (B). A gradient elution method was employed, starting from 75% A and ending at 53% A over 60 minutes, with a flow rate of 0.35 mL min^−1^. Prior to injection, 25 μL of purified procainamide-labeled *N*-glycans were mixed with 75 μL of ACN to optimize loading conditions. A 40 μL portion of this mixture was injected for analysis. Instrument control was managed using Hyster 4.1 software (Bruker Daltonik, GmHB). MS conditions included a capillary voltage of 4.5 kV, a source temperature of 250 °C, nebulizer gas set at 1.7 bar, and drying gas flow at 6 L min^−1^. MS spectra were acquired within the range of 50 to 2800 m/z at a frequency of 1 Hz. MS/MS experiments targeted the two most abundant precursor ions, with spectra rates ranging from 0.5 Hz to 2 Hz. Collision energies varied depending on precursor charge states, with specific values assigned for doubly, triply, and quadruply charged precursors. Stepping-energy experiments utilized a basic stepping mode with collision RF values set at 1500 and 2100 Vpp (each for 50% of the time).

### 2.6 Data analysis

The identification of procainamide-labeled *N*-glycans was conducted using Protein Scape software V4 (Bruker Daltonik, GmHB). Initially, data from the HILIC-FLD-QTOF-MS/MS analysis was converted to .xml file format using Data Analysis software (Bruker Daltonik, GmbH). Converted data was then processed within Protein Scape. To determine *N*-glycan structures, the tandem mass spectra of procainamide-labeled N-glycans was searched against the GlycoQuest Search Engine. The search parameters included MS and MS/MS tolerances set to 20 ppm and 0.05 Da, respectively, with CarbBank as the database (Doubet & Albersheim, 1992). A threshold score of 10 was applied to identify the procainamide-labeled *N*-glycans. In addition, extracted ion chromatograms of the m/z ratio of precursors were generated, and their structures were subsequently annotated manually. Glycoworkbench was used for the illustration of *N*-glycan structures (Ceroni et al., 2008).

### 2.7 Diversity and data analyses

Glycan relative abundances were used to calculate the Shannon Entropy (Shannon, 1948) and the number of observed features, as well as the Bray Curtis distance metric as a measure of beta diversity (Bray & Curtis, 1957). Bray Curtis distances were compared using an adonis PERMANOVA test for group significance (Anderson, 2001; Dixon, 2003). Alpha diversity results were compared using the nonparametric Kruskal-Wallis test (Kruskal & Wallis, 1952).

## 3. Results

### 3.1 Elucidated N-glycan structures across glycoproteins

HPLC-HILIC-FLD-QTOF-MS/MS analysis revealed a total of 33, 33, 10, and 10 *N-*glycan structures from bovine whey, egg, soy, and pea protein isolates, respectively. Interestingly, the *N-*glycans produced from animal sources generally had a greater number of distinct *N-*glycans when compared to the plant sources, with 33 for both bovine whey and egg white and only 10 distinct *N-*glycans for both soy and pea proteins. The most abundant *N-* glycan per protein source were distinct compared to each other, shown in Figure 1. Table 1 shows a list of all the *N-*glycans represented by all four chromatograms with their *m/z* values, charges, relative abundances, and their glycoprotein sources.

**Figure 1.**
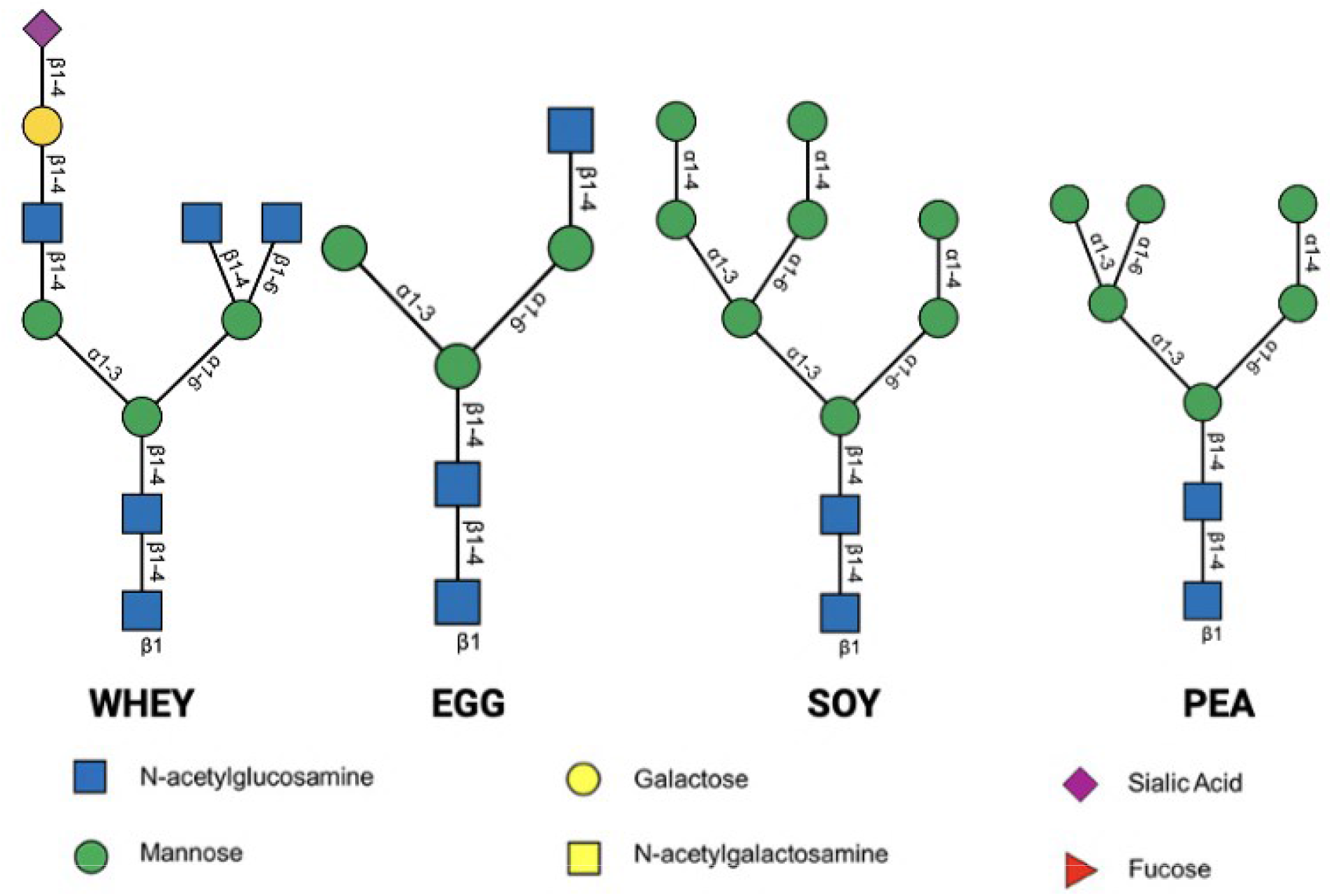
The most abundant *N-*glycan structures, as determined by mass spectrometry. The most abundant *N*-glycan per glycoprotein source is shown, as measured by HPLC paired with QTOF mass spectrometry to identify *N-*glycan pools from four distinct dietary glycoproteins.

QTOF-MS/MS identified 22 peaks, which resulted in 33 total *N-*glycan structures for bovine whey. Bovine whey contained the most diverse *N-*glycans by monosaccharide composition with 5 distinct sugars, including *N*-acetylglucosamine (GlcNAc), mannose, galactose, fucose, and N-acetylneuraminic acid (Neu5Ac) (Figure 2). Bovine whey protein was the only protein that contained acidic *N-*glycans, with a total of 12 acidic structures out of the 33 total structures identified. Bovine whey also contained *N-*glycans decorated with fucose monosaccharides unlike the other protein sources, which did not contain any fucose decorations. The most abundant *N-*glycan represented 23.28% of the total *N-* glycome of bovine whey, with the second and third most abundant *N-*glycans at 11.92% and 8.05%, respectively.

**Figure 2.**
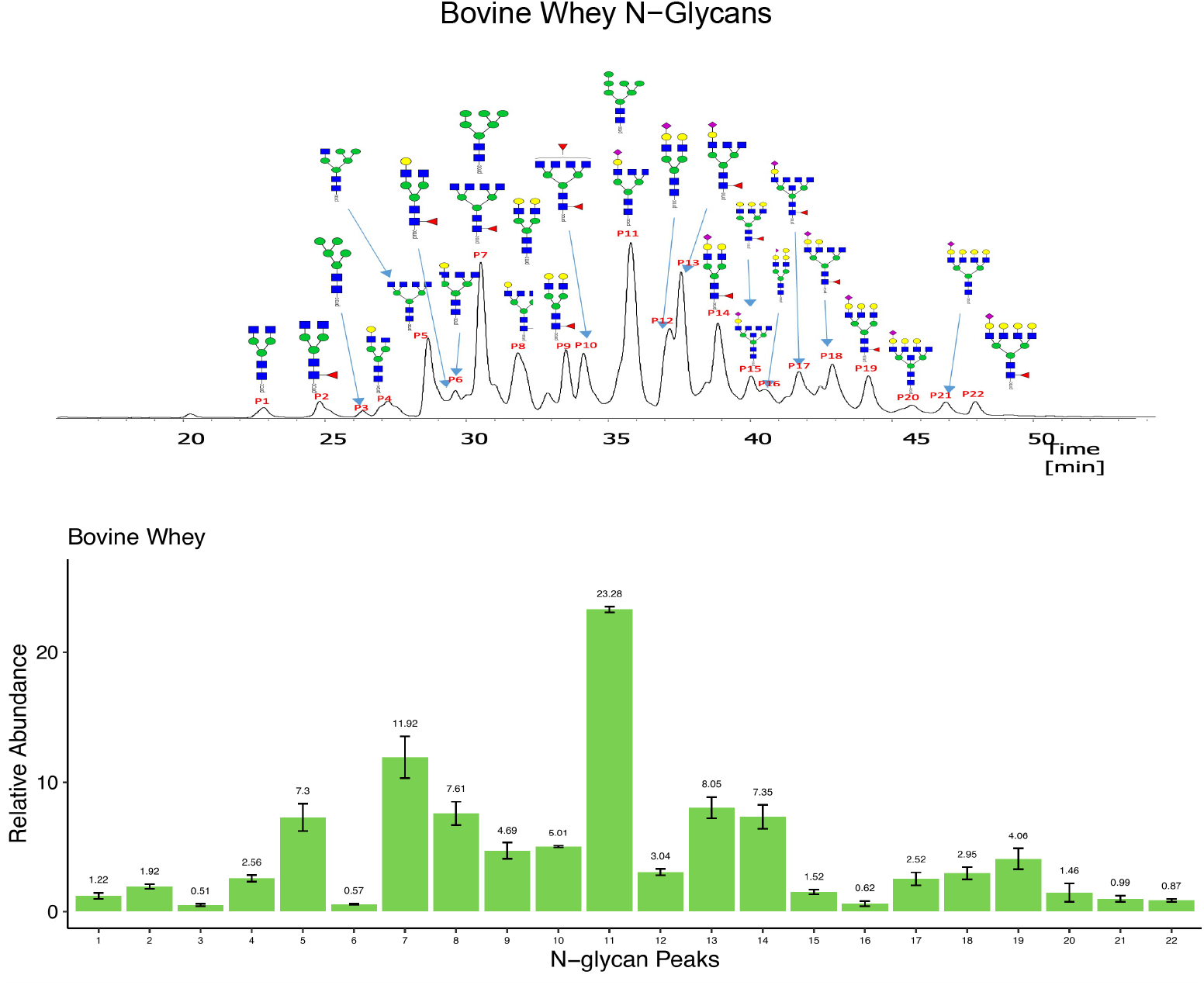
Distinct *N-*glycan structures from bovine whey glycoprotein and their relative abundance determined by QTOF-MS/MS. HILIC-HPLC with a fluorescence detector paired with QTOF-MS/MS produced chromatograms then determined structures and abundance for procainamide-labeled *N*-glycans from bovine whey protein. A total of 22 peaks were identified corresponding to 33 distinct *N-*glycan structures. *N-*glycan structures for low abundance peaks not shown.

With similarities to the bovine whey protein in terms of the number of distinct *N-*glycan structures, QTOF-MS/MS identified 30 peaks representing 33 *N-*glycan structures for egg white protein. The monosaccharide composition decorating the egg white *N-*glycome was compromised of *N*-acetylglucosamine, *N*-acetylgalactosamine, mannose, and galactose with no acidic *N-*glycans present (Figure 3). Interestingly, *N*-acetylgalactosamine was unique to the *N-*glycome of egg white protein. The three most abundant egg white *N-*glycans were nearly evenly distributed at 12.72%, 9.84%, and 9.65%, respectively.

**Figure 3.**
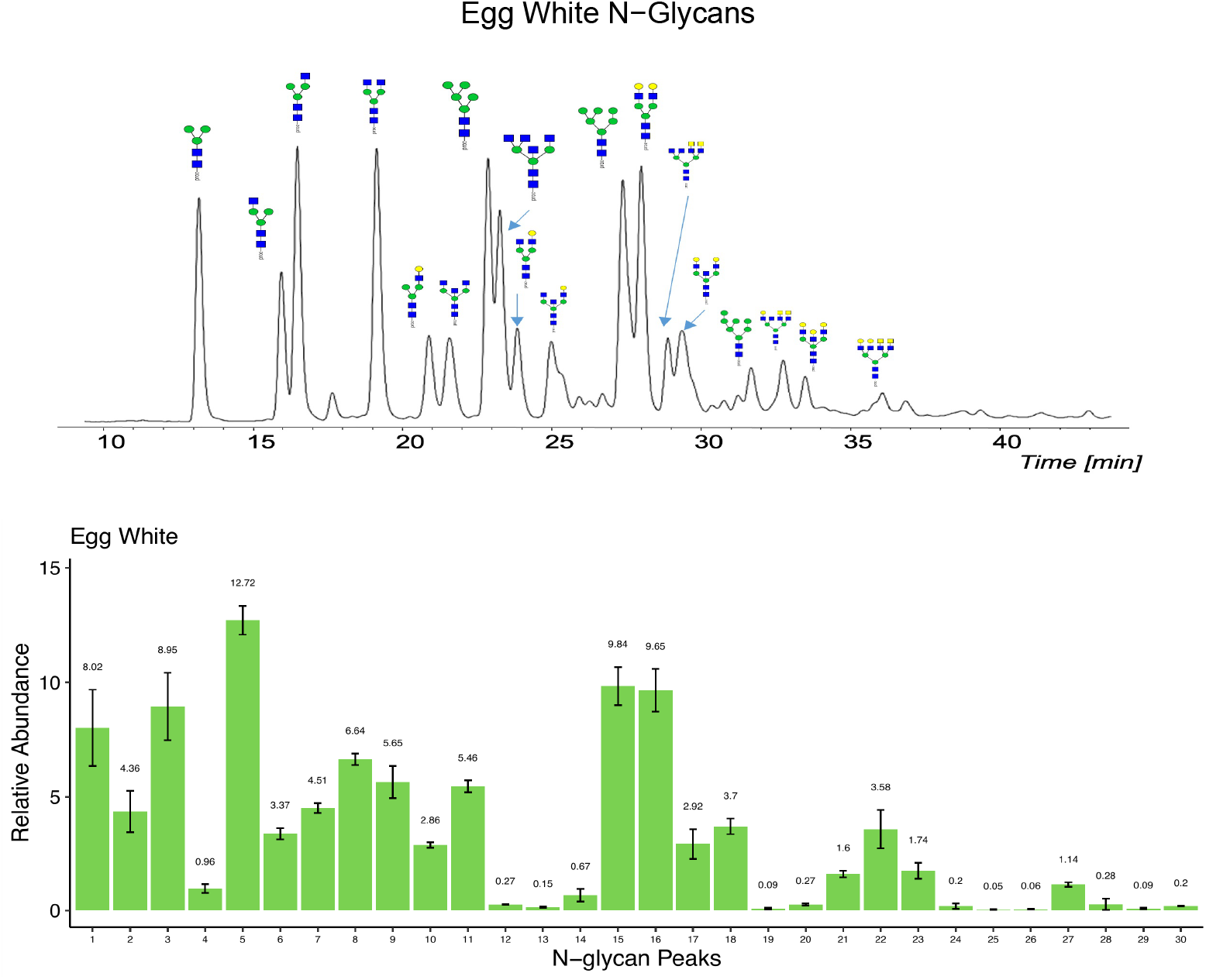
Distinct *N-*glycan structures from egg white glycoprotein and their relative abundance determined by QTOF-MS/MS. HILIC-HPLC with a fluorescence detector paired with QTOF-MS/MS produced chromatograms then determined structures and abundance for procainamide-labeled *N*-glycans from egg white protein. A total of 30 peaks were identified corresponding to 33 distinct *N-*glycan structures. *N-*glycan structures for low abundance peaks not shown.

Unlike both animal protein sources, soy protein contained 10 peaks representing 10 distinct *N-*glycans which were predominantly oligomannosidic structures (Figure 4). Unique to soy protein, 60.69% of the *N-*glycome was represented by the most abundant *N-*glycan. The second and third most abundant *N-* glycans made up 23.14% and 11.48%, respectively. These top 3 most abundant structures made up over 90% of soy’s *N-*glycome.

**Figure 4.**
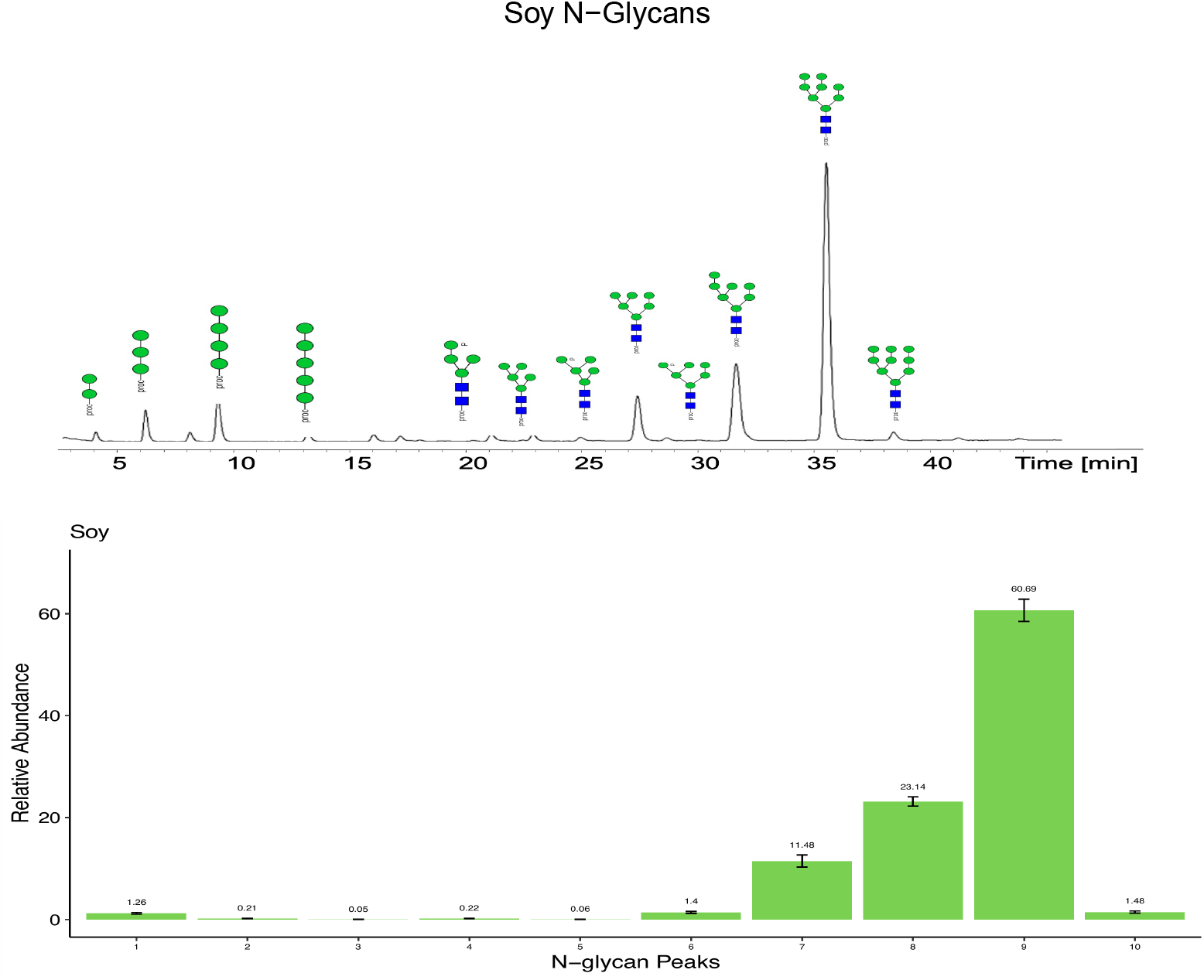
Distinct *N-*glycan structures from soy glycoprotein and their relative abundance determined by QTOF-MS/MS. HILIC-HPLC with a fluorescence detector paired with QTOF-MS/MS produced chromatograms and determined structures and abundance for procainamide-labeled *N*-glycans from soy protein. A total of 10 peaks were identified corresponding to 10 distinct *N-*glycan structures. *N-*glycan structures for low abundance peaks not shown.

Similar to the soy *N-*glycome, QTOF-MS identified 10 peaks corresponding to 10 distinct *N-*glycans that were predominantly oligomannosidic which decorated pea protein (Figure 5). Pea protein contained a more even distribution of *N-*glycans compared to soy protein, with the most abundant structures representing 39.25% of the *N-*glycome. The second and third most abundant structures represented 24.43% and 17.66% of the *N-*glycome, respectively.

**Figure 5.**
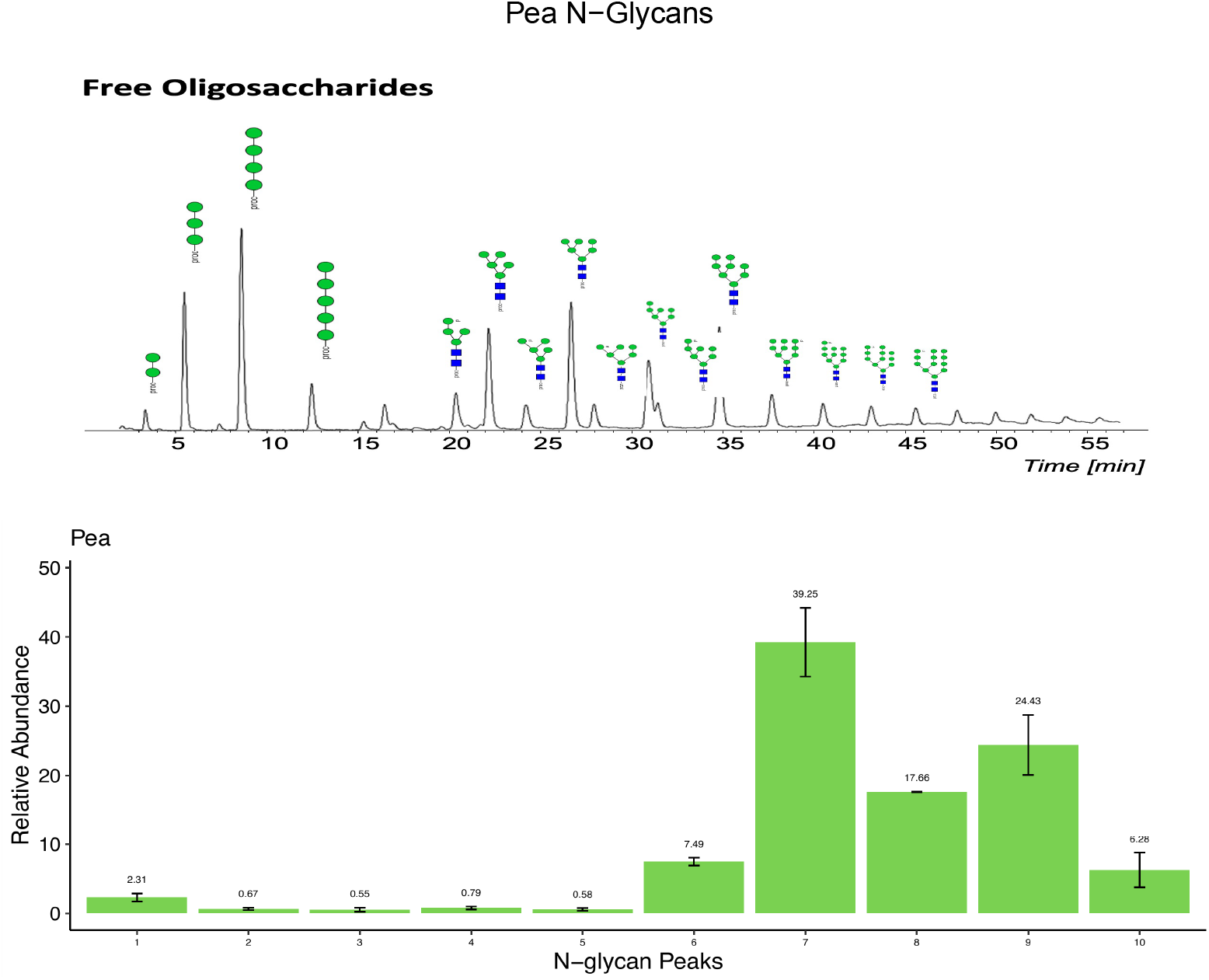
Distinct *N-*glycan structures from pea glycoprotein and their relative abundance determined by QTOF-MS/MS. HILIC-HPLC with a fluorescence detector paired with QTOF-MS/MS produced chromatograms and determined structures and abundance for procainamide-labeled *N*-glycans from pea protein. A total of 10 peaks were identified corresponding to 10 distinct *N-*glycan structures. *N-*glycan structures for low abundance peaks not shown.

### 3.2 N-glycome comparison between animal and plant sources

Consistent with the literature, we found stark differences in *N-*glycan monosaccharide composition, or *N-*glycan “type” between the animal and plant sources. While the animal sources contained oligomannosidic *N-*glycans, the predominant *N-* glycan type found within both sources were complex *N-*glycans composed of multiple monosaccharide types. This is in contrast to the plant sources’ *N-*glycomes, which were exclusively oligomannosidic *N-* glycan types. Interestingly, both animal sources contained roughly 20 additional structures compared to the plant sources, revealing the increased *N-*glycan diversity among the animal sources compared to the plant sources. Among all protein sources, bovine whey glycoprotein shared 11 (21%) *N-*glycan structures with egg white protein and 5 (10%) with both soy and pea protein, and included 20 (38%) unique structures (Figure 6A). The pea and soy *N-*glycome included 10 structures, which were shared (Figure 6A), though the abundance of each *N-*glycan differed between the plant proteins (Figure 4, 5). Finally, the egg white *N-*glycome shared 4 (8%) structures with both the pea and soy *N-*glycomes, while containing 14 (27%) unique *N-*glycan structural compositions (Figure 6A), which included 7 pairs of structural isomers (Table 1). 3 (6%) *N-*glycan structures were shared among all glycoprotein sources (Figure 6A). When comparing animal- and plant-derived *N-*glycans, 6 *N-*glycan structures (12%) were shared between both plant and animal sources, with 42 (81%) and 4 (8%) unique *N-*glycan structures found within the animal sources and plant sources, respectively (Figure 6B).

**Figure 6.**
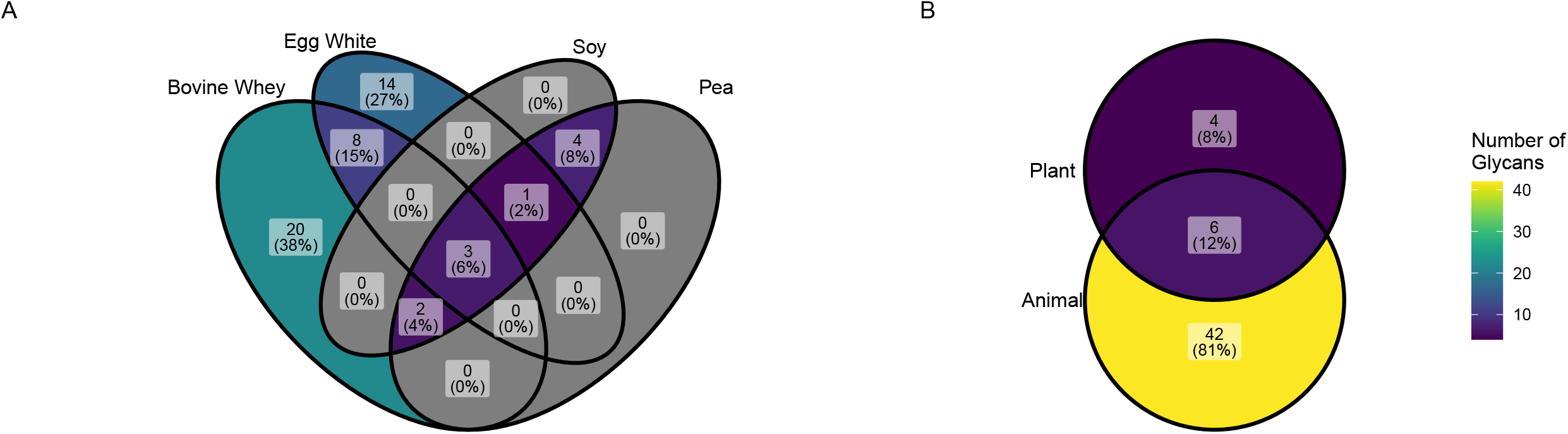
Some *N-*glycan structures are unique, and others are shared between glycoproteins and glycoprotein sources determined by HPLC-QTOF-MS/MS. (A) Comparison of *N-*glycan structures between all glycoprotein sources determined by HPLC-QTOF-MS/MS. (B) Comparison of *N-* glycan structures between animal and plant sources.

### 3.3 Glycan diversity and similarity

When the number of structures and their distribution among samples were compared, egg white and whey protein had the most diverse *N*-glycans by both diversity measures (Figure S1A). Egg white and whey protein had a mean Shannon entropy of 4.31 (+/- 0.024 SD) and 4.34 (+/- 0.059 SD) respectively, while pea and soy protein had a mean Shannon entropy of 2.26 (+/-0.039 SD) and 1.59 (+/- 0.064 SD), respectively. For both observed features and Shannon entropy measures, the differences between group comparisons were significant (P < 0.001 and P < 0.005, respectively). The β-diversity comparisons for these *N-*glycoproteins also demonstrated significant differences when compared by a PERMANOVA test (P = 0.001).

When the protein sources were grouped by their source type (e.g., animal vs. plant), the protein sources were also significantly different. Both the β-diversity measure and α-diversity measures (Shannon entropy and Observed features) were significantly different (P = 0.002, P < 0.005, and P < 0.001, respectively; Figure S1B).

## 4. Discussion

Here, we used HPLC-HILIC-FLD-QTOF-MS/MS to analyze four distinct and widely consumed sources of dietary glycoproteins from phylogenetically diverse and commercially important sources. These protein sources represent four of the most abundant sources of dietary protein among both whole and processed foods, as well as supplemental dietary protein (Smeuninx et al., 2020). We characterized the structural differences in *N-*glycan structures between these glycoprotein sources. HPLC-HILIC-FID-QTOF-MS/MS analysis was used to ensure proper separation, purification, and identification of complex pools of *N-*glycans.

In all, a total of 33, 33, 10, and 10 *N-*glycan structures were identified from bovine whey, egg, soy, and pea protein isolates, respectively. The main *N*-glycome difference across protein sources is found in the composition and arrangement of monosaccharides and the number of distinct *N*-glycans per protein source. Soy and pea proteins are primarily decorated with oligomannosidic *N-*glycans, whereas egg and bovine whey proteins possess structures with a wider variety of monosaccharides (**Figure 1**). *N-* glycans derived from egg protein included mannose, galactose, *N*-acetylglucosamine, and *N-* acetylgalactosamine while bovine whey included the most complex and diverse *N-*glycans, decorated with mannose, galactose, *N*-acetylglucosamine, sialic acid, and fucose. Given the findings in terms of the number of distinct glycans, it is unsurprising that the diversity of structures among *N-*glycoproteins differed primarily between animal and plant-derived glycoproteins. Soy and pea glycoproteins, and their composition of primarily oligomannose structures, lacked the structural diversity that was observed among the egg white and whey proteins. These findings reflect the evolutionary origins of protein *N*-glycosylation and empirical comparisons between glycoproteins in the plant and animal kingdoms (Pedrazzini et al., 2016; Wang et al., 2017).

The determination of the *N-*glycome for each of four protein ingredients identified consistent findings in *N-*glycan architecture congruent with a broader understanding of *N-*glycan biosynthesis. Previous studies examining the *N-*glycome of bovine whey have consistently identified complex structures containing sialic acids and fucose (Nwosu et al., 2012; Valk-Weeber, Deelman-Driessen, et al., 2020; Valk-Weeber, Eshuis-de Ruiter, et al., 2020; van Leeuwen et al., 2012), which is consistent with our findings. Other work that has characterized the *N-*glycome for soy protein allergens are also in agreement with our findings of primarily oligomannosidic *N-*glycan structures (Li et al., 2016), with some limited incorporation of monosaccharides such as xylose and *N-*glycan core fucosylation (Lu et al., 2022; Zhu et al., 2018). These minor differences can most likely attributed to differences in methodologies. The identification of the egg white *N-*glycome is more limited, with most studies examining the egg *N-*glycome focusing on species that are not widely regarded as food sources (Sanes et al., 2019; Suzuki et al., 2004, 2009), or they have examined the egg yolk *N-*glycome from different animals (Kayili, 2021; Roth et al., 2010). To our knowledge, this is the first report of the hen egg white and pea *N-*glycomes.

One limitation of the present work is that the method used here is not as sensitive as other methods to elucidate *N-*glycan structures, such as matrix-assisted laser desorption/ionization mass spectrometry (MALDI-MS) and it is difficult to confidently annotate structures for the lesser peaks in the MS spectra. However, HPLC-HILIC-FID-QTOF-MS/MS provides more consistent data for semi-quantitative analysis of *N*-glycans and is able to confidently identify bisecting *N*-glycan structures based on the procainamide labeling approach.^15^ Further, we have not examined how and whether genetic or environmental determinants across breeds or strains of livestock or crops affects *N*-glycosylation, though there is evidence that free oligosaccharides and endogenous glycosidases found in bovine milk can vary,(Robinson et al., 2019; Sunds et al., 2021) and the *N*-glycome of human milk is subject to variation.(Barboza et al., 2012; Smilowitz et al., 2013) Thus while there may be additional and unappreciated variation beyond the present work, when these protein sources are used in food manufacturing, their inclusion is in a form comparable to the substrates examined here.

The identification of the *N-*glycome of dietary glycoprotein sources is an important contribution to the developing study of how *N-*glycan composition can shape the gut microbiome. It is likely that the observed differences in the complexity and decoration of *N-*glycans impact the human gut microbiome in different ways, however the distinct impact of glycoprotein amino acid sequences and *N-*glycan composition remains to be determined.

## 7. Figure Legends

**Supplementary Figure 1.**
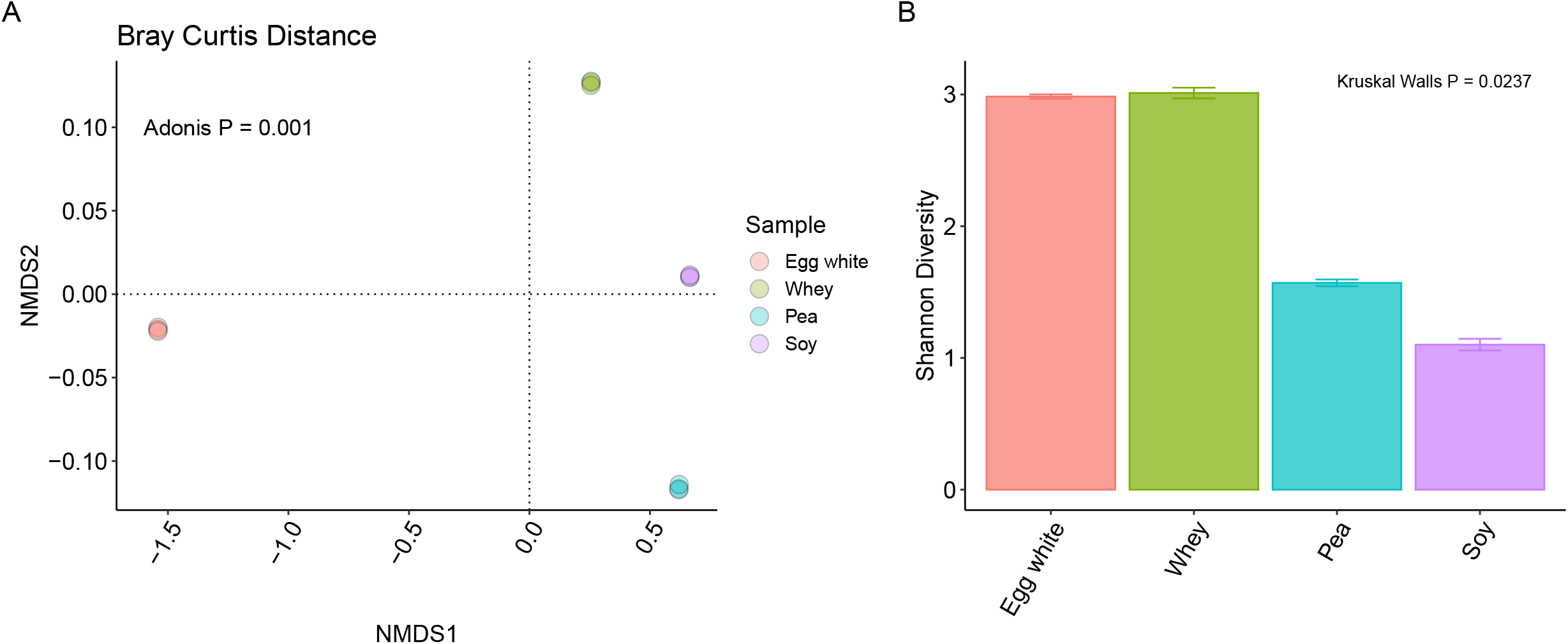
Alpha- and beta-diversity of *N-*glycans from glycoprotein sources determined by HPLC-QTOF-MS/MS. (A) A Bray-Curtis dissimilarly metric was used to compare *N-* glycans pools across glycoproteins, which identified significant differences between these *N*-glycan pools (Adonis P = 0.001). (B) The Shannon diversity, or alpha diversity, of *N-*glycan pools per glycoprotein is shown. A Kruskal Walls test identified significant differences in Shannon diversity across glycoprotein sources (P = 0.0237).

